# Development of a viability real time PCR assay for Bordetella pertussis, B parapertussis and B holmesii

**DOI:** 10.1101/2025.11.04.686548

**Authors:** Kalichamy Alagarasu, Savita Katendra, Meera Prabhakar, Rajlakshmi Viswanathan

**Author notes:** **Corresponding author:** Rajlakshmi Viswanathan, ICMR-National Institute of Virology, 20-A. Dr Ambedkar Road Pune 411001, Maharashtra, India, /26006890. Email IDs: Kalichamy Alagarasu.

## Abstract

**Background**

Real time PCR provides rapid and accurate laboratory confirmation of clinically suspected pertussis and pertussis like illness (PLI). However, it does not inform the viability as it detects bacterial DNA from both viable and degraded bacterial cells. We report development of platinum based viability real time PCR assays for *Bordetella pertussis, B*.*parapertussis* and *B*.*holmesii* which cause pertussis and PLI.

**Material and Methods:** ATCC strains of the three organisms were grown on selective media, and cell counts determined. Untreated (control) and Sodium dodecyl sulphate (SDS) lysed cultures were treated with varying concentrations of platinum chloride (PtCl_4_) and genomic DNA extracted. Real time PCR was performed for the targets of *B*.*pertussis (IS481), B*.*parapertussis* (*pIS1001*) and *B*.*holmesii* (*hIS1001)* respectively, and quantitative results obtained. threshold cycle values after PtCl_4_ exposure and before PtCl_4_ exposure (Δ Cq) were calculated for both untreated and SDS treated cultures.

**Results:** For all three species, increase in ΔCq values was noted with corresponding reduction in DNA copy numbers, as PtCl_4_ concentration was increased, in untreated and SDS treated cultures. Untreated cultures of all three species exposed to PtCl_4_showed increase in Cq values, suggesting that liquid cultures contain a mixture of live and dead bacteria.

**Conclusion:** Performance of viability q PCR assay incorporating PtCl_4_ at a concentration range of 6 to 10 mM would inform the viability of the infecting organism in the clinical sample. This assay would improve the diagnostic reliability, help select samples for culture as well as adoption of mitigation strategies among contacts of pertussis cases.

**Importance:**

- Pertussis and pertussis like illness (PLI) are highly contagious respiratory tract infections for which Real time PCR is the frontline diagnostic tool.
- This test however does not inform the viability of the infecting pathogen, as it detects nucleic acid which can be present even after degradation of the cell.
- Performing a viability qPCR incorporating platinum chloride (PtCl_4_) at a concentration range of 6 to 10 mM would inform the viability of *B. pertussis, B. parapertussis* and *B. holmesii* in the clinical sample.
- It is critical to know the viability of the infecting organism to determine which cases of pertussis and PLI which are actually infectious, to adopt mitigation strategies including prophylactic antibiotics among close contacts
- A viability assay can be used for quantitative measurement of bacterial number and viability and will help select the correct specimens for culture and isolation of the pathogen.

## Introduction

*Bordetella pertussis* is the causative agent of pertussis, commonly known as whooping cough. The disease usually presents as a prolonged afebrile or minimally febrile cough illness but the clinical features may vary depending on the age or immunization status. Pertussis like illness (PLI) is also caused by other species of *Bordetella* like *B*.*holmesii* and *B*.*parapertussis* (Bouchez and Guiso, 2015; Njamkepo et al., 2011). Laboratory testing is therefore crucial for confirmation of etiology. The gold standard for laboratory confirmation of pertussis is culture of nasopharyngeal swabs or aspirates. However, it is laborious, requires special media and has a long turnaround time. Real time PCR (qPCR) provides a rapid and accurate diagnosis, and is increasingly in use for confirmation of pertussis and other *Bordetella* spp (Martini et al., 2017; Tatti et al., 2011). The assay however does not inform the viability of the infecting pathogen, as the test detects nucleic acid which can be present even after degradation of the cell. The viability of the organism is critical for determining the infectivity of the patient. Pertussis is a highly contagious disease and known to spread to family and other close contacts (Sali et al., 2015). It is important to know if the patient having a prolonged cough illness of 3-4 weeks, continues to remains a source of infection. Viability assays are also important for human challenge studies (Ramkissoon et al., 2020).

A qPCR assay using propidium monoazide (PMA) was reported, to enumerate live and dead cells of *B*.*pertussis* (Ramkissoon et al., 2020).The method requires special conditions like incubation in the dark and use of a halogen lamp for irradiation. An alternative is the use of platinum compounds which have been effective in viability PCR assays for contaminating pathogens like *E*.*coli* and *Cronobacter sakazakii* in milk (Soejima et al., 2016), hepatitis A and E viruses in environmental water samples (Randazzo et al., 2018), noroviruses (Fraisse et al., 2018) and for differentiation of viable and non-viable SARS-CoV-2 (Cuevas-Ferrando et al., 2021). Platinum compounds are known to be chelated by nucleic acid ligands in mammalian cells and are usually used in cancer chemotherapy (Lovejoy et al., 2008). They can penetrate non-viable cells with a compromised cell surface. The platinum compounds interact with nucleic acid like DNA resulting in cross-links, which interfere with the amplification of the target nucleic acid sequence in qPCR. Therefore, when detecting microbial viability, only unmodified DNA from live cells is amplified (Soejima et al., 2016). They also have the advantage of not being light sensitive or requiring excitation by light to function (Soejima et al., 2016).

We developed a quantitative real time PCR assay for differentiation of viable and non-viable *B. pertussis* using platinum chloride. We also developed similar assays for *B*.*holmesii* and *B. parapertussis*, which are known to cause pertussis like illness. These assays can be used effectively in both diagnostics and research studies where a quantitative measure of, and information on viability of these three species of *Bordetella* is required.

## Methods

### Bacterial strains, culture and viable cell count

*B*.*pertussis* (ATCC 9797) was inoculated in Brucella broth (Hi Media Laboratories, Mumbai,India), *B. parapertussis* (ATCC 15311) in Bordet Gengou broth (Hi Media Laboratories, Mumbai, India), and *B. holmesii* (ATCC 51541) in brain heart infusion broth (BD, BBL, Sparks Maryland, USA). The cultures were incubated at 37^0^C for 72 hours for *B*.*pertussis* and *B. holmesii*, and at 35°C with 5% CO_2_ for *B. parapertussis*.

For *B*.*pertussis*, the plate grown colonies were resuspended in 1 mL Brucella broth to an OD_600_ = 1.0 (approximately 10^8^ cfu/ml) for estimation of cell count as described earlier (Li et al., 2014; Ramkissoon et al., 2020). For *B*.*parapertussis* and *B*.*holmesii*, the viable cell counts were determined by standard plate count method (Murray et al., 2004) on Bordet Gengou Agar (Hi Media Laboratories, Mumbai, India) and sheep blood agar (Hi Media Laboratories, Mumbai, India) respectively, with tenfold dilutions of the cultures prepared in 1 ml PBS using the formula

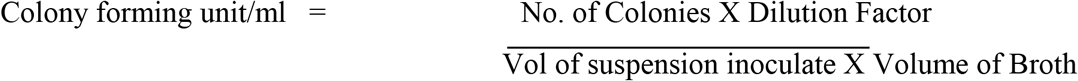

The culture plates were incubated upto 72 hours at 37°C for *B. holmesii* and 35°C with 5% CO_2_ for *B. parapertussis*.

#### Preparation of SDS treated working stock of bacterial cultures

To simulate dead bacterial cells which would be present in clinical samples, the live bacterial cultures of the three *Bordetella* spp, were treated with sodium dodecyl sulfate (SDS) to prepare working stocks. In brief, 1.5 ml of a 1% solution of SDS (Biomol, Hamburg, Germany) in 0.2 N NaOH was mixed with equal volume of broth culture of each of the three bacterial strains. The suspension was intermittently vortexed for 20 minutes and used for further assays. To confirm bacterial death after treatment, 10 µl of the suspension of each of the three strains was inoculated as described above, on solid culture media and incubated up to seven days to observe for bacterial growth (Birnboim and Doly, 1979; Wada et al., 2012).

#### Preparation of Platinum chloride (PtCl_**4**_)

PtCl_4_ (Sigma-Aldrich, Saint Louis, MO, United States) was dissolved in dimethyl sulfoxide (DMSO) (Sigma-Aldrich, Saint Louis, MO, United States) at a concentration of 300 millimolar (mM) to prepare a master stock solution, which was stored at -20°C. This was further diluted in phosphate buffered saline (PBS) to prepare a 100mM working solution, which was also stored at -20°C until further use.

#### Exposure of untreated and treated bacterial cells to PtCl_**4**_

The untreated and SDS treated bacterial cells were exposed to different concentrations of PtCl_4_. Hundred microliter each of the suspensions of *B. pertussis, B. parapertusis* and *B. holmesii* were separately mixed with different volumes of 100 mM PtCl_4_ (Table 1). A final volume of 1000 µl was made up with PBS, to attain a final concentration of 2mM, 4mM, 6mM, 8Mm and 10 mM of PtCl_4_. The suspensions were well-mixed and incubated on rotor at 22°C to 25°C for 30 minutes.

**Table 1:** Preparation of bacterial suspensions with different concentration of PtCl_4_.

| Volume of 100mM PtCl<sub>4</sub> solution (μL) | 20 | 40 | 60 | 80 | 100 |
| --- | --- | --- | --- | --- | --- |
| PBS (μL) | 880 | 860 | 840 | 820 | 800 |
| Bacterial Culture (μL) | 100 | 100 | 100 | 100 | 100 |
| Final Concentration (mM) of PtCl<sub>4</sub> | 2 | 4 | 6 | 8 | 10 |

#### Extraction of Genomic DNA

After incubation, both the live cultures of the three *Bordetella* spp. (controls), as well as SDS treated bacterial suspensions exposed to PtCl_4_, were centrifuged at 14000 rpm for three minutes. The pellets were re suspended in the buffer provided with the commercial extraction kit to make up the final volume of 180 µl for DNA extraction using the Qiagen DNA Mini (Qiagen, DNA Mini kit, USA) as per manufacturer’s instructions. The extracted DNA was eluted using 50 µl of elution buffer.

#### Standard curve and sensitivity of qPCR Assay

Quantitation standard curves were established by real time PCR (Tatti et al., 2011) on the Bio-Rad CFX96 (Bio-Rad laboratories, Singapore), using ten 10-fold dilutions of known concentration of the genomic DNA of the three species of *Bordetella*. The copy number per reaction based on known genomic DNA concentration was calculated using the formula

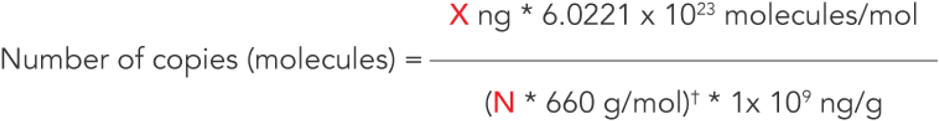

Copy number per millilitre was further determined using the formula, Copies/µl X elution volume(µl)/sample volume(ml)

The standards were tested in duplicates in each run and on three consecutive days.

#### Taqman Real time PCR Assay

Single target qPCR assays were used to detect target DNA from viable bacterial cells of *B*.*pertussis, B*.*parapertussis* and *B*.*holmesii* in the live culture (controls) and SDS treated suspensions exposed to different concentrations of PtCl_4_, The assays in triplicate, were performed on a Bio-Rad CFX 96 (Bio-Rad laboratories, Singapore) for detection of target genes for *B*.*pertussis* (*IS481), B parapertussis* (*pIS1001*), and *B holmesii (hIS1001*) using Taqman Gene expression Master mix (Thermo Scientific, Waltham, MA USA) as described earlier (Tatti et al., 2011). A cycle threshold value (Cq) of ≤ 35 cycles was considered for positivity. Cq values >35 and < 40 were considered indeterminate. Any Cq value ≥40 was considered as not detected. A negative control and one extraction control were included in each qPCR reaction.

DNA concentration and the length of template values were used to calculate the copy number per microliter. Δ Cq values i.e. Cq (after PtCl_4_ exposure) - Cq (before PtCl_4_ exposure) were calculated for both untreated and SDS treated bacterial cultures.

## Results

### Cell count

The cell count for *B. pertussis* cell culture OD_600_ was 1.388=1.1*10^9^cell/ml. The viable cell count was 1.1*10^9^ cfu/ml and 3*10^8^cfu/ml for *B. parapertussis* and *B. holmesii* respectively. The three bacterial suspensions did not show any growth after treatment with SDS.

### Quantification of Platinum treated bacterial suspensions and controls by q PCR Assay

The mean Cq values for different copy number of DNA/ml are provided in supplementary table 1 and the standard curves for calculating the copy number of DNA of the thee pathogens are shown in figure 2. The results for untreated and SDS treated bacterial suspensions exposed to PtCl_4_ for the three species of *Bordetella* is presented in Table 2. The threshold cycle value (Cq) and calculated copy number per ml for live cultures of *B. pertussis, B*.*parapertussis and B*.*holmesii* was 9.39 (33.95*10^12^), 18.28(70.5*10^12^) and 15.03 (53.23*10^12^) respectively. Similarly for the SDS treated cultures of *B*.*pertussis, B*.*parapertussis* and *B*.*holmesii*, the Cq and calculated copy number per ml was 21.94(33.95*10^8^), 16.27(7.05*10^10.5^), and 18.46(53.23*10^9.5^) respectively.

**Table 2:**
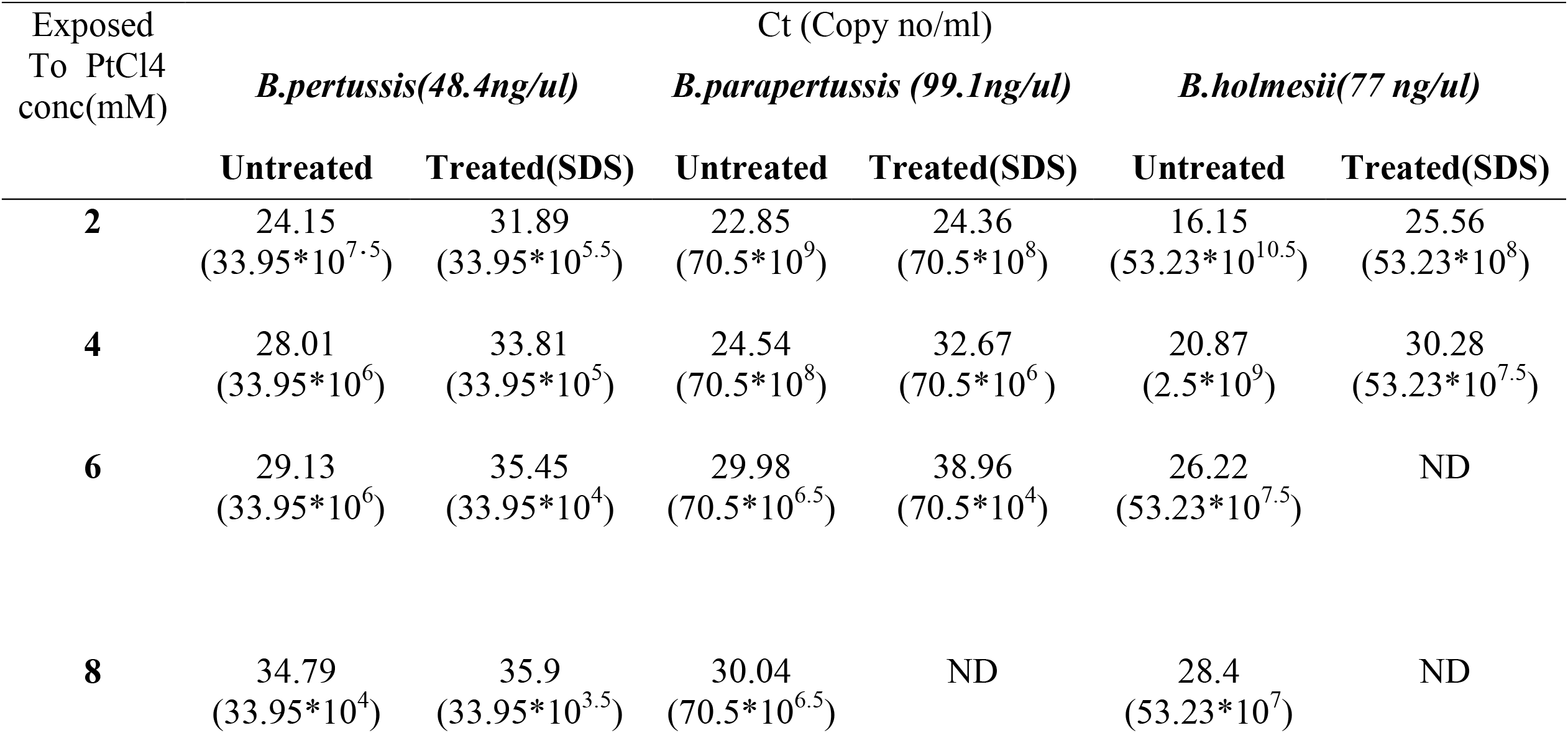
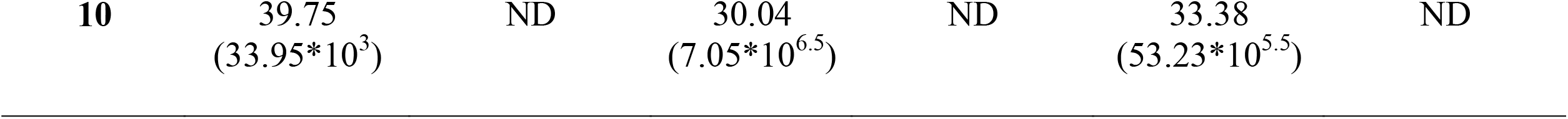
Live and SDS treated of three *Bordetella* spp exposed to different concentrations of PtCl_4_.

**Fig 1.**
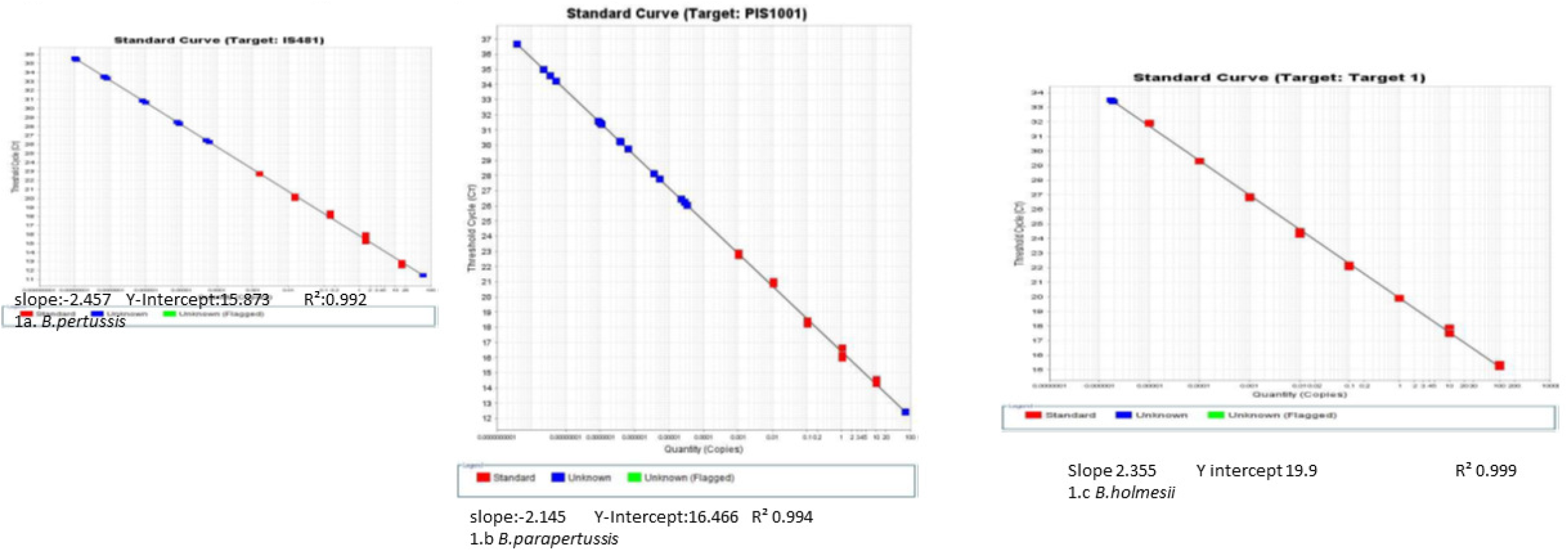
Standard curves for Different copy number of DNA/ml from *Bordetel la pertussis, B*.*parapertussisand B*.*holmessi:*

**Fig 2.**
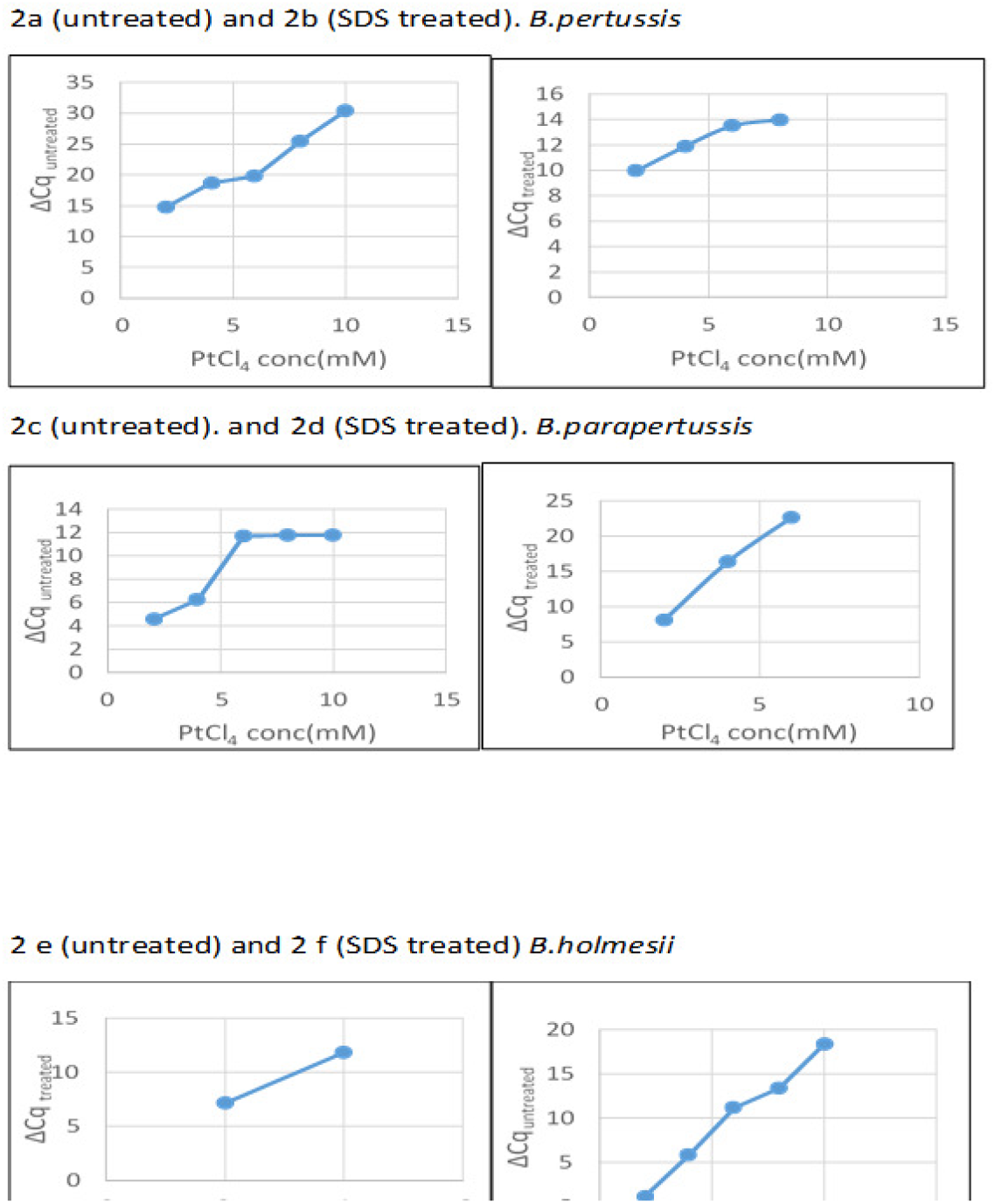
Δ Cq for untreated and treated bacterial suspensions 2a (untreated) and 2b (SDS treated). *a*.*pertussis*

For all three *Bordetella* species, there was increase in ΔCq values with increase in concentration of PtCl_4_, both in live and SDS treated cultures (Table 3). For live cultures of *B*.*pertussis*, amplification was detected upto 10mM concentration of PtCl4 with positivity upto 8mM PtCl_4_ concentration (Fig2a and b). Whereas, for live cultures of *B*.*parapertussis* (Fig 2 c and d) and *B*.*holmesii* (Fig 2 e and f), positivity was detected upto the maximum concentration of PtCL_4_ used (10mM).

**Table 3:**
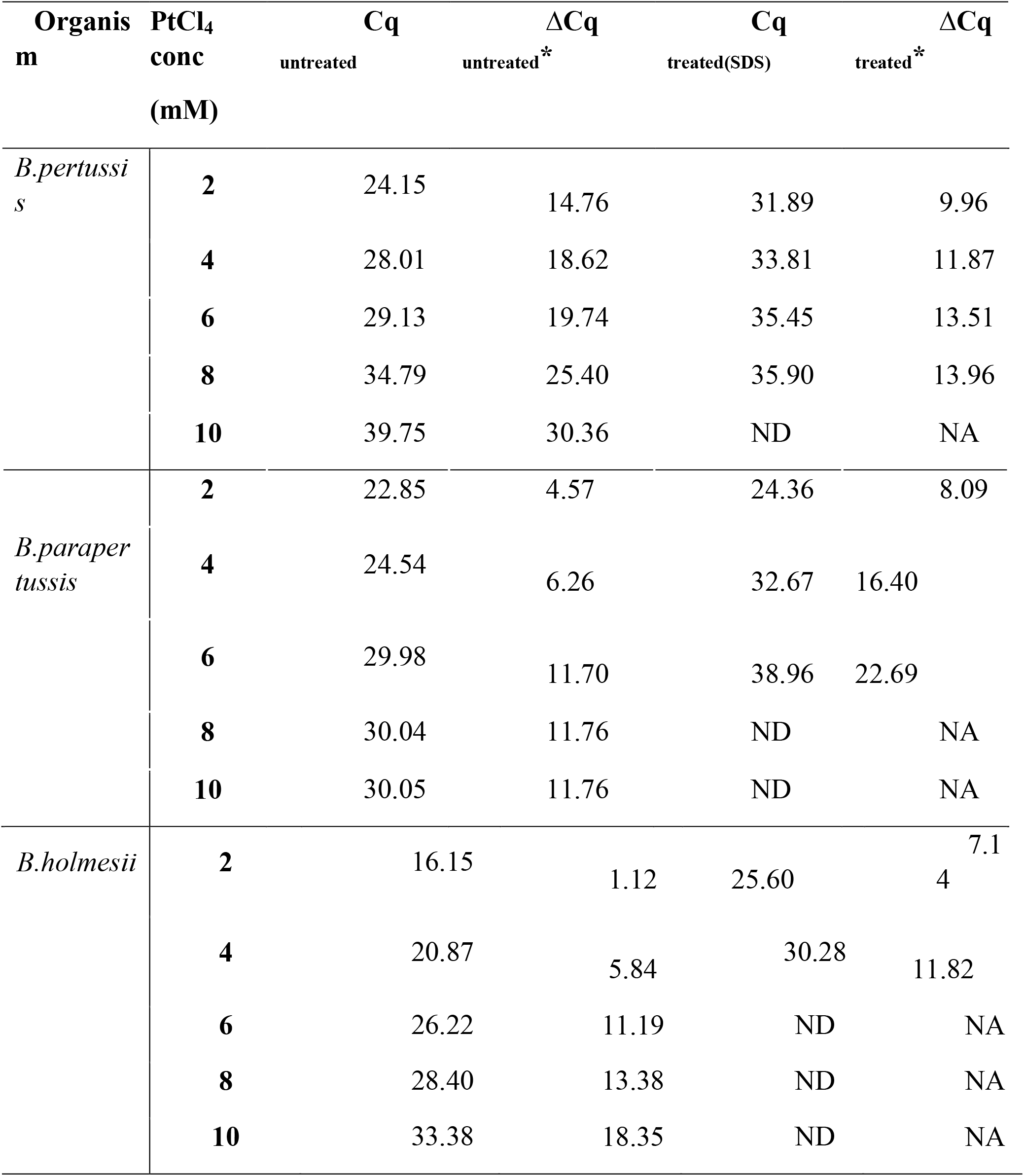
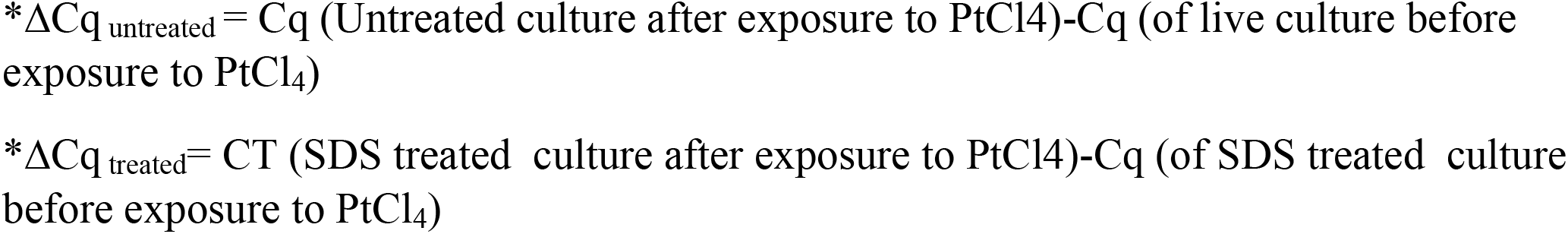
ΔCq for untreated and treated bacterial cultures of three *Bordetella*.

| Organism | PtCl<sub>4</sub><br>conc<br>(mM) | Cq | ΔCq | Cq | ΔCq |
| --- | --- | --- | --- | --- | --- |
|  |  | untreated | untreated* | treated(SDS) | treated* |
| *B.pertussis* | 2 | 24.15 | 14.76 | 31.89 | 9.96 |
|  | 4 | 28.01 | 18.62 | 33.81 | 11.87 |
|  | 6 | 29.13 | 19.74 | 35.45 | 13.51 |
|  | 8 | 34.79 | 25.40 | 35.90 | 13.96 |
|  | 10 | 39.75 | 30.36 | ND | NA |
| *B.paraper*<br>*tussis* | 2 | 22.85 | 4.57 | 24.36 | 8.09 |
|  | 4 | 24.54 | 6.26 | 32.67 | 16.40 |
|  | 6 | 29.98 | 11.70 | 38.96 | 22.69 |
|  | 8 | 30.04 | 11.76 | ND | NA |
|  | 10 | 30.05 | 11.76 | ND | NA |
| *B.holmesii* | 2 | 16.15 | 1.12 | 25.60 | 7.1<br>4 |
|  | 4 | 20.87 | 5.84 | 30.28 | 11.82 |
|  | 6 | 26.22 | 11.19 | ND | NA |
|  | 8 | 28.40 | 13.38 | ND | NA |
|  | 10 | 33.38 | 18.35 | ND | NA |

\* $\Delta Cq_{\text{untreated}} = Cq$ (Untreated culture after exposure to $PtCl_4$ )- $Cq$ (of live culture before exposure to $PtCl_4$ )
\* $\Delta Cq_{\text{treated}} = CT$ (SDS treated culture after exposure to $PtCl_4$ )- $Cq$ (of SDS treated culture before exposure to $PtCl_4$ )

When pre-treatment with SDS was done, there was increase in ΔCq values with increase in concentration of PtCl_4_ upto 6 mM for *B*.*pertussis* which plateaued and was similar at 8mM. At 10mM concentration, amplification was completely suppressed. (Fig2a and b).

For *B*.*parapertussis* results were negative at 6mM concentration of PtCl_4_ and amplification was completely suppressed beyond this concentration of PtCl_4_,(Fig 2 c and d) whereas for *B*.*holmesii*, amplification was completely suppressed at 6mM concentration of PtCl_4_ itself (Fig 2 e and f),

On exposure to different concentrations of PtCl_4,_ the untreated cultures of all three *Bordetella* spp. showed increase in Cq values (Table 3), suggesting that the liquid cultures actually contain a mixture of live and dead bacteria. While positivity was detected in case of *B*.*parapertussis and B*.*holmesii* upto 10mM concentration of PtCl_4_, in case of *B*.*pertussis* this was only upto 8mM, and the results were similar at this concentration to the SDS treated cultures.

## Discussion

Pertussis and pertussis like illness are highly contagious respiratory tract infections, particularly in children. With a resurgence of pertussis in recent years, there is an urgent need to improve and expand the diagnostic tools for these infections. One such tool would be the development of assays for providing information to undertake preventive strategies and countermeasures for these infections.

We undertook the development of platinum based viability PCR assays for *B*.*pertussis, B*.*parapertussis* and *B*.*holmesii*. The assays would enable the identification of the cases of pertussis and PLI which are actually infectious. A viability PCR for *B*.*pertussis* using PMA has been reported (Ramkissoon et al., 2020). However as mentioned earlier, this reagent is light sensitive and also requires additional equipment like a halogen lamp for activation, which are not needed for platinum compound based viability assays. An earlier study (Soejima et al., 2016) has evaluated the use of several platinum compounds for bacterial viability PCR assays. PtCl_4_ which was identified as the most effective platinum compound for distinguishing the viability of bacterial cells was used in our study. To simulate the presence of dead bacteria in clinical samples, the cultures of three *Bordetella* spp. were pre-treated with an alkaline solution of SDS, a widely used bactericidal surfactant (Yilmaz and Icgen, 2014), which is considered as a typical cell wall perturbing agent and used routinely for lysis of Gram negative organisms (Wada et al., 2012).

Viability PCR assays for bacteria have correlated the result of the qPCR assay with reduction of infectivity in terms of cfu/ml (Soejima et al., 2016). The assay has used a target which is an integral part of the organism, like 16srRNA for Enterobacteriacae (Soejima et al., 2016). Insertion sequences (IS) are the most widely accepted diagnostic target for detection of *B. pertussis, B*.*parapertussis* and *B*.*holmesii* and have been been used in our study (Tatti et al., 2011). The use of the same target as in the standard diagnostic algorithm will facilitate the implementation of the viability q PCR assay in laboratories. These targets are sensitive but present in varying copy numbers even within the same species and as such cannot be directly correlated with the colony count in cfu/ml (Tatti et al., 2011). A PMA based viability PCR assay for pertussis using the same target as our study (Ramkissoon et al., 2020) has also not correlated the results with colony count in cfu/ml.

Our results for untreated and SDS treated cultures of *B*.*pertussis,B*.*parapertussis* and *B*.*holmesii*, show that as PtCl_4_ concentration is increased, there is increase in Cq values. Performance of a viability q PCR assay incorporating PtCl_4_ at a concentration range of 6 to 10 mM would inform the viability of the infecting organism in the clinical sample.

Our data suggest that even the untreated cultures of all three *Bordetella* spp, which showed increase in Cq values on exposure to PtCl_4,_ actually contain a mixture of live and dead bacteria. A limitation of our study was that further investigation to enumerate the live and dead bacteria could not be carried out. *B*.*pertussis* is a relatively slow growing fastidious organism which means the usual methods of plate count for quantification cannot be undertaken. Even if performed, the results may not be accurate as the solid culture may contain a mixture of live and dead bacteria (Ramkissoon et al., 2020). Our data suggest that cultures of *B*.*parapertussis* and *B*.*holmesii* on solid media may also actually contain a similar mixture of live and dead bacteria.

Therefore the adoption of PtCl_4_ based q PCR assays, can help to accurately distinguish viable from non-viable bacterial cells, making it a promising tool for understanding the viability of *B*.*pertussis* and other species causing PLI. These assays would help to improve the diagnostic reliability and minimise false-positive results. The laboratory culture of *B*.*pertussis* is challenging due to its fastidious growth requirements. The capability of PtCl_4_ to differentiate viable and non-viable bacterial cells will help to identify samples which need to be taken up for culture. If the result of PtCl_4 -_qPCR assay indicates the predominance of non-viable cells, the need for culture can be eliminated. In conclusion, the results of this viability PCR assay will help to inform strategies including prophylactic antibiotics for mitigating the risk of infection among contacts of pertussis cases.

## Data Statement

The data presented in this study are available on request from the corresponding author.

## Author Contribution

**KA** Conceptualization, Supervision, Reviewing and Editing. **MP, SK:** Methodology, Data curation. **RV:** Conceptualization, Supervision Writing-Original draft preparation, Reviewing and Editing,project administration, funding acquisition. All authors have read and agreed to the published version of the manuscript.

## Institutional Review Board Statement

The study was conducted according to the guidelines of the Declaration of Helsinki, and approved by the Institutional Ethics Committee of ICMR-National Institute of Virology (protocol code 18-1-3 and 24th June 2019).

## Acknowledgments

Prof. Veeraraghavan Balaji, Department of Clinical Microbiology,Christian Medical College, Vellore for sharing real time PCR protocols for *Bordetella pertussis* and other species,.

---

**Supplementary Table 1:**
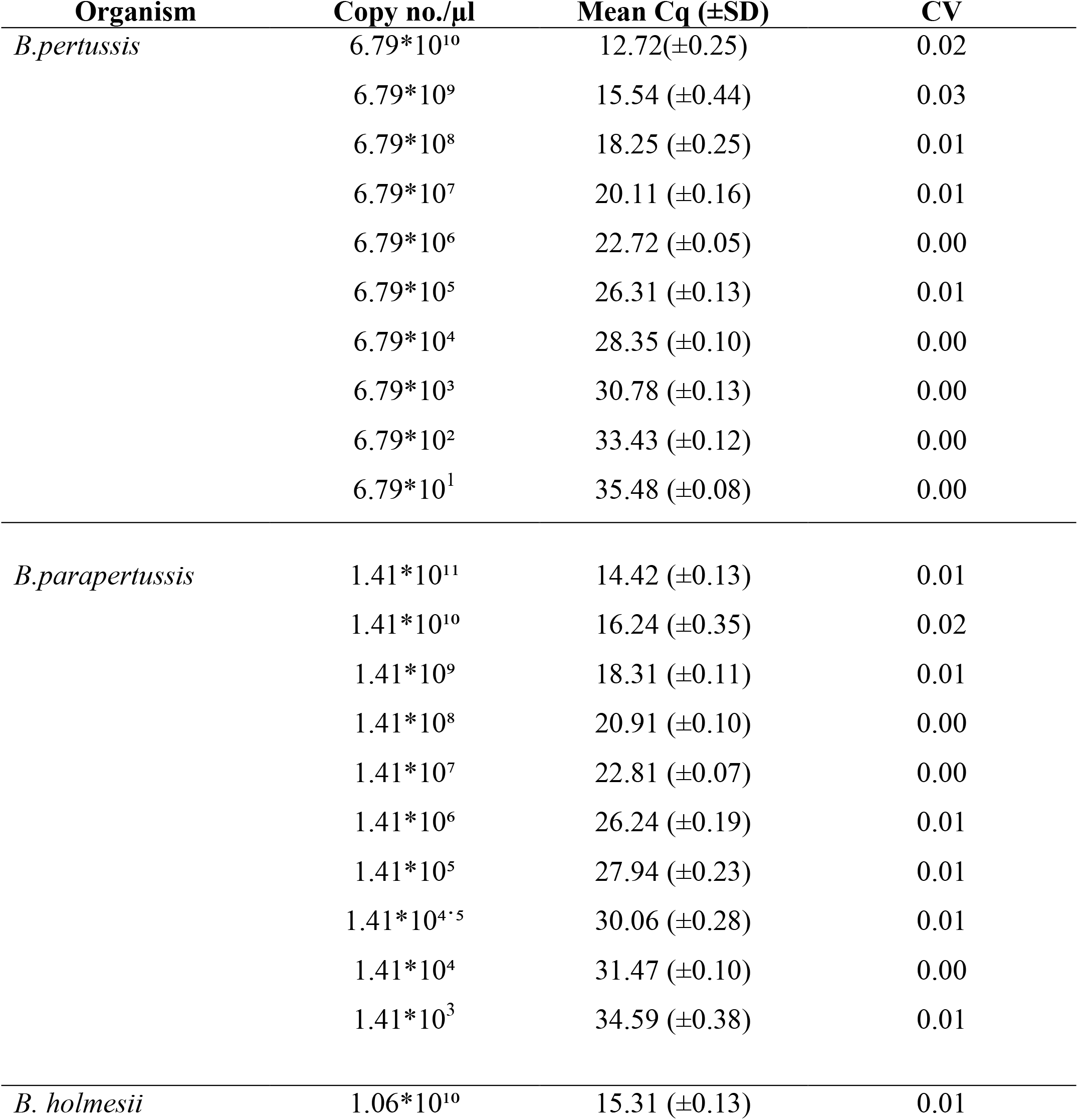
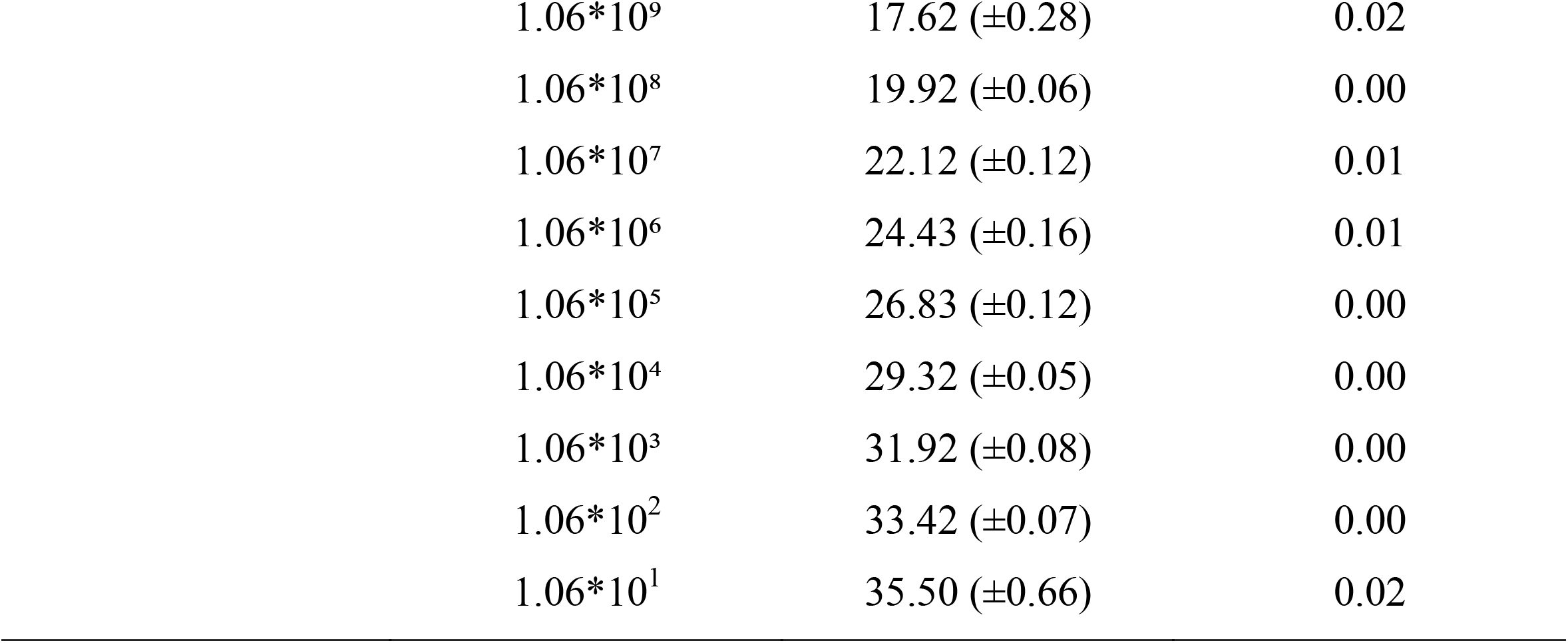
Number of DNA Copies number per microliter with mean Cq values of three *Bordetella* spp.

| Organism | Copy no./$\mu l$ | Mean $Cq$ ($\pm SD$) | CV |
| --- | --- | --- | --- |
| *B.pertussis* | $6.79 \times 10^{10}$ | 12.72($\pm 0.25$) | 0.02 |
| | $6.79 \times 10^9$ | 15.54 ($\pm 0.44$) | 0.03 |
| | $6.79 \times 10^8$ | 18.25 ($\pm 0.25$) | 0.01 |
| | $6.79 \times 10^7$ | 20.11 ($\pm 0.16$) | 0.01 |
| | $6.79 \times 10^6$ | 22.72 ($\pm 0.05$) | 0.00 |
| | $6.79 \times 10^5$ | 26.31 ($\pm 0.13$) | 0.01 |
| | $6.79 \times 10^4$ | 28.35 ($\pm 0.10$) | 0.00 |
| | $6.79 \times 10^3$ | 30.78 ($\pm 0.13$) | 0.00 |
| | $6.79 \times 10^2$ | 33.43 ($\pm 0.12$) | 0.00 |
| | $6.79 \times 10^1$ | 35.48 ($\pm 0.08$) | 0.00 |
| *B.parapertussis* | $1.41 \times 10^{11}$ | 14.42 ($\pm 0.13$) | 0.01 |
| | $1.41 \times 10^{10}$ | 16.24 ($\pm 0.35$) | 0.02 |
| | $1.41 \times 10^9$ | 18.31 ($\pm 0.11$) | 0.01 |
| | $1.41 \times 10^8$ | 20.91 ($\pm 0.10$) | 0.00 |
| | $1.41 \times 10^7$ | 22.81 ($\pm 0.07$) | 0.00 |
| | $1.41 \times 10^6$ | 26.24 ($\pm 0.19$) | 0.01 |
| | $1.41 \times 10^5$ | 27.94 ($\pm 0.23$) | 0.01 |
| | $1.41 \times 10^{4.5}$ | 30.06 ($\pm 0.28$) | 0.01 |
| | $1.41 \times 10^4$ | 31.47 ($\pm 0.10$) | 0.00 |
| | $1.41 \times 10^3$ | 34.59 ($\pm 0.38$) | 0.01 |
| *B. holmesii* | $1.06 \times 10^{10}$ | 15.31 ($\pm 0.13$) | 0.01 |

|  |  |  |
| --- | --- | --- |
| $1.06 \cdot 10^9$ | $17.62 (\pm 0.28)$ | 0.02 |
| $1.06 \cdot 10^8$ | $19.92 (\pm 0.06)$ | 0.00 |
| $1.06 \cdot 10^7$ | $22.12 (\pm 0.12)$ | 0.01 |
| $1.06 \cdot 10^6$ | $24.43 (\pm 0.16)$ | 0.01 |
| $1.06 \cdot 10^5$ | $26.83 (\pm 0.12)$ | 0.00 |
| $1.06 \cdot 10^4$ | $29.32 (\pm 0.05)$ | 0.00 |
| $1.06 \cdot 10^3$ | $31.92 (\pm 0.08)$ | 0.00 |
| $1.06 \cdot 10^2$ | $33.42 (\pm 0.07)$ | 0.00 |
| $1.06 \cdot 10^1$ | $35.50 (\pm 0.66)$ | 0.02 |

